# Urolithin A Restores Mitochondrial Function and Reverses Cardiac Remodeling in Heart Failure with Preserved Ejection Fraction

**DOI:** 10.64898/2025.12.26.696570

**Authors:** Hangyul Song, Chahyun Yun, Yun Ju Choi, Wooju Jeong, Yumin Kim, Jaeyoung Kim, Ju Yeon Lee, Dongryeol Ryu, Sang-Wook Park, Chang-Myung Oh

## Abstract

**Background:** Heart failure with preserved ejection fraction (HFpEF) accounts for nearly half of all heart failure cases; however, mechanism-based therapies targeting cellular dysfunction remain absent. Emerging evidence suggests that mitochondrial dysfunction and impaired quality control, particularly defective mitophagy, are central to the pathogenesis of HFpEF. Urolithin A (UA), a gut microbiome-derived postbiotic metabolite, has shown promise as a mitophagy activator in preclinical models; however, its therapeutic efficacy in HFpEF remains unknown.

**Methods:** We used a clinically relevant two-hit HFpEF mouse model (high-fat diet (HFD) plus Nω-nitro-L-arginine methyl ester (L-NAME)) and co-administered UA during disease progression. The cardiac structure and function were assessed using echocardiography, histology, and transmission electron microscopy. Mitochondrial bioenergetics were evaluated using Seahorse-based respirometry. Mitophagy flux was monitored using mt-Keima assays in H9c2 cells and human-induced pluripotent stem cell-derived cardiomyocytes (hiPSC-CMs). Mechanistic insights were obtained using immunoblotting, integrated shotgun metagenomic and lipidomic profiling, and single-nucleus RNA sequencing (snRNA-seq).

**Results:** UA treatment significantly attenuated cardiac remodeling and fibrosis in HFpEF mice while improving diastolic function parameters. Transmission electron microscopy revealed restoration of the mitochondrial ultrastructure, and mitochondrial stress tests demonstrated enhanced oxidative phosphorylation capacity and glycolytic reserves. Immunoblot assays revealed that UA treatment recovered PINK1/Parkin-mediated mitophagy markers (reduced LC3-II and p62/SQSTM1). Mt-Keima assays confirmed enhanced mitophagic flux in H9c2 cells under HFpEF-like stress conditions. Integrated metagenomics and lipidomics revealed significant reductions in cardiovascular risk-associated ceramides. snRNA-seq demonstrated that HFpEF-like stress downregulated contractile, calcium-handling, and mitochondrial mitophagy gene programs. UA treatment restored the mitochondrial quality control signatures and normalized profibrotic and conduction-type cell populations.

**Conclusions:** Urolithin A restored mitochondrial quality control through the activation of mitophagy and reversed cardiac remodeling in HFpEF. These findings establish UA as a mitochondria-targeted therapeutic candidate for HFpEF.

**Novelty and Significance:** *What is known?:*

✓ Heart failure with preserved ejection fraction (HFpEF) is a heterogeneous syndrome driven by metabolic stress and myocardial remodeling, for which effective disease-modifying therapies are lacking.
✓ Mitochondrial dysfunction and impaired mitophagy have been implicated in HFpEF, but their roles in cardiomyocyte state remodeling and fibrosis remain incompletely defined.
✓ Urolithin A, a gut microbiome–derived metabolite, enhances mitophagy in aging and metabolic tissues, but its relevance to HFpEF has not been established.

*What new information does this article contribute?:*

✓ Urolithin A restores mitochondrial quality control and mitophagic flux in cardiomyocytes, improving diastolic function and attenuating cardiac remodeling in a two-hit HFpEF model.
✓ Integrated metagenomic and lipidomic analyses identify suppression of gut microbiome–associated ceramide biosynthesis as a systemic mechanism linked to Urolithin A–mediated cardioprotection.
✓ Single-nucleus RNA sequencing reveals that Urolithin A reverses maladaptive cardiomyocyte transcriptional state transitions toward fibrogenic and conduction-like programs.

## INTRODUCTION

Heart failure with preserved ejection fraction (HFpEF) accounts for approximately half of all heart failure cases and represents a major unmet clinical need worldwide^1^. HFpEF is a heterogeneous syndrome driven by multiple comorbidities, including obesity, hypertension, and diabetes, which converge to induce systemic inflammation, endothelial dysfunction, and myocardial remodeling. Although sodium-glucose cotransporter-2 inhibitors have recently shown modest clinical benefits, mechanism-based therapies addressing the fundamental cellular pathology of HFpEF remain absent^2,3^.

Mitochondrial dysfunction has emerged as a central pathogenic mechanism in HFpEF^4,5^. The enormous energy demand of the heart necessitates efficient oxidative phosphorylation and robust mitochondrial quality control (MQC) to sustain the contractile and relaxation functions. In HFpEF, this homeostasis is disrupted, leading to the accumulation of damaged mitochondria, which generate excessive reactive oxygen species, impair ATP production, and dysregulate calcium handling^4,5^. A critical component of MQC is mitophagy, the selective autophagic removal of dysfunctional organelles^6^. Defective mitophagy has been implicated in age-related cardiovascular diseases and metabolic cardiomyopathy, suggesting that the restoration of this clearance mechanism represents a rational therapeutic strategy^6–8^. However, no clinically approved mitophagy-enhancing therapies exist for HFpEF, and whether such approaches can reverse the established pathology of this complex syndrome remains to be determined.

Urolithin A (UA) is a gut microbiome-derived metabolite produced from dietary ellagitannins found in pomegranates, berries, and walnuts^9^. UA activates mitophagy across species ranging from *C. elegans* to humans, improving muscle function and extending the healthspan^10–12^. Although preclinical studies have demonstrated cardioprotective effects in ischemia-reperfusion and diabetic cardiomyopathy models^13,14^ the therapeutic efficacy of UA in HFpEF, where metabolic stress and structural remodeling predominate, has not yet been investigated. Here, we investigated the therapeutic potential of UA in a two-hit mouse model of HFpEF that combined a high-fat diet and nitric oxide synthase inhibition to recapitulate the key metabolic and hemodynamic features of the human disease^15,16^. We hypothesized that UA would ameliorate diastolic dysfunction by restoring mitophagy and improving mitochondrial homeostasis. Using an integrated multiomics approach encompassing shotgun metagenomics, plasma lipidomics, and single-nucleus RNA sequencing, we delineated systemic and cell-autonomous mechanisms linking mitochondrial restoration to the suppression of profibrotic and inflammatory programs. Our findings demonstrated that UA ameliorated HFpEF pathology in this preclinical model through the coordinated modulation of mitophagy, ceramide metabolism, and fibrogenic programs and provided a mechanistic rationale for its therapeutic development in HFpEF.

## METHODS

### 2.1 Cell cultures

H9c2 cardiomyocytes were maintained in Dulbecco’s Modified Eagle’s Medium (DMEM) supplemented with 10% fetal bovine serum FBS and exposed to Control, HF-like stress (palmitate+L-NAME), or HF+UA conditions for 24 h before downstream molecular, imaging, and functional analyses, as indicated^17–19^. Human iPSCs were cultured on Matrigel and differentiated into cardiomyocytes using a chemically defined cardiac induction protocol. Beating began on days 8–10, and the experiments were performed on days 15–20. Minus-insulin selection was used when enrichment was required^20–22^.

### 2.2 Mouse experiments

All animal experiments were approved by the Institutional Animal Care and Use Committee of Gwangju Institute of Science and Technology (GIST-2021-110) and conducted in accordance with NIH and ARRIVE guidelines. Eight-week-old male C57BL/6J mice were randomly assigned to Control, HFpEF, or HFpEF+UA groups. HFpEF was induced using a two-hit protocol consisting of a high-fat diet (60% kcal from fat) and L-NAME administration in drinking water, whereas Control mice received standard chow and water. UA was administered by mixing into the high-fat diet at indicated doses. Body weight was monitored weekly. Cardiac function and body composition were assessed by transthoracic echocardiography and dual-energy X-ray absorptiometry (DXA), respectively.^23–26^.

### 2.3 Echocardiography

Transthoracic echocardiography was performed using small-animal ultrasound systems (VINNO 6 LAB or VisualSonics Vevo 2100) under isoflurane anesthesia. Left ventricular systolic function was evaluated from midventricular short-axis M-mode images, including ejection fraction and fractional shortening. Diastolic function was assessed from apical four-chamber views using pulsed-wave and tissue Doppler imaging to measure transmitral inflow parameters (E and A waves). Measurements were performed according to established guidelines^27–29^.

### 2.4 Histological analysis

Hearts were fixed in 4% paraformaldehyde, paraffin-embedded, and sectioned at 6 μm. Cardiac morphology and fibrosis were assessed by hematoxylin and eosin and Masson’s trichrome staining respectively. Fibrotic area was quantified using ImageJ/Fiji under blinded conditions^30–32^. For ultrastructural analysis, fixed heart tissues were processed for transmission electron microscopy. Mitochondrial number, size, and structural integrity were quantified from randomly selected fields^30–32^.

### 2.5 Transmission Electron Microscopy (TEM)

Ultrastructural analysis of mitochondrial morphology was performed on fixed heart tissues. Ultrathin sections were imaged at high magnification, and the mitochondrial number, size, and structural integrity were quantified from multiple randomly selected fields per sample^33–35^.

### 2.6 Immunoblotting & qRT-PCR

Protein extracts from cells or tissues were subjected to SDS-PAGE and immunoblotting for autophagy/mitophagy and signaling markers, including ULK1, PINK1, Parkin, LC3, p62, AMPK, mechanistic target of rapamycin (mTOR), and AKT. Band intensities were quantified by densitometry and normalized to β-actin. Total RNA was isolated, reverse-transcribed, and analyzed by quantitative real-time PCR. Relative mRNA expression levels were calculated using the ΔΔCt method^36^ and normalized to housekeeping genes. Primer sequences are provided in Supplementary Table 1.36

### 2.8 Bulk sequencing (RNA-seq)

Sequencing data were processed using Python-based pipelines to obtain normalized expression values. Differentially expressed genes were identified using Wald tests, with effect sizes summarized as log₂ fold changes. P values were integrated using Stouffer’s method, followed by false discovery rate correction. Genes meeting significance criteria were subjected to Gene Ontology and Gene Set Enrichment Analysis^37,38,39^.

### 2.9 Lipidomics

Plasma lipidomic profiling was conducted using the MxP Quant 500XL kit (Biocrates Life Sciences AG). Metabolites were quantified by LC–MS/MS and flow injection analysis, and data were processed using the Biocrates MetIDQ platform with internal quality controls applied. Metabolites failing quality criteria were excluded from downstream analyses. Additional details are provided in the Supplementary Methods^40^.

### 2.10 Shotgun metagenomic sequencing

Fecal DNA was extracted and subjected to shotgun metagenomic sequencing. After quality filtering and host-read removal, taxonomic and functional profiling were performed using MetaPhlAn v3.0 and HUMAnN v3.0, respectively, with gene families regrouped to KEGG orthologs. Alpha diversity was assessed using the Shannon index, whereas beta diversity was quantified using Bray-Curtis dissimilarity. Additional details are provided in the Supplementary Methods^41,42,43^.

### 2.11 Mito Stress assay & mitophagy assays

Mitochondrial respiration was assessed using the Seahorse XF Mito Stress Test. Basal respiration, ATP-linked respiration, maximal respiration, and spare respiratory capacity were calculated, with oxygen consumption rates normalized to total protein content^44^. Mitophagy was evaluated using the mt-Keima assay in H9c2 cardiomyocytes. Ratiometric fluorescence imaging was performed under identical acquisition settings, and mitophagy was quantified as the per-cell 561/458 nm fluorescence ratio. Carbonyl Cyanide m-Chlorophenyl Hydrazone (CCCP) served as a positive control.

### 2.13 Single-nucleus RNA sequencing

Single-nucleus RNA sequencing was performed on hiPSC-derived cardiomyocytes using a 10x Genomics platform. Data were normalized, integrated, clustered, and visualized by UMAP. Differential expression analyses were performed using single-nucleus and pseudobulk approaches, and cardiomyocyte state transitions were inferred using pseudotime trajectory analysis implemented in scFates. Detailed procedures are described in the Supplementary Methods.

### 2.14 Statistical analysis

Data are presented as mean ± standard error of the mean (SEM). Group comparisons were performed using two-way ANOVA with Dunnett’s post hoc test or Kruskal–Wallis test with Dunn’s correction, as appropriate. Transcriptomic analyses used Wald tests with false discovery rate correction, and p values from multiple tests were combined using Stouffer’s method. Statistical significance was defined as P<0.05 or FDR<0.05.

## RESULTS

### 3.1 UA treatment restored diastolic function in a two-hit HFpEF mouse model

To model the complex pathophysiology of HFpEF, 8-week-old male C57BL/6J mice were subjected to a two-hit protocol that combined high fat diet (HFD) and L-NAME administration for 8 weeks (Fig. 1A). Echocardiography at week 8 confirmed successful induction of the HFpEF phenotype; the ejection fraction remained within the normal range, yet the diastolic function was markedly impaired, as demonstrated by a significantly elevated E/A ratio, indicative of restrictive filling (Fig. 1B). After phenotypic confirmation, the mice continued the two-hit regimen with the co-administration of UA for an additional 12 weeks. Notably, UA treatment had no effect on the body weight trajectories, daily food consumption, or body composition parameters, including lean and fat masses (Figs. 1C, D), indicating that the metabolic effects of the HFpEF-inducing regimen were not altered by UA and suggesting metabolic tolerability.

**Figure 1.**
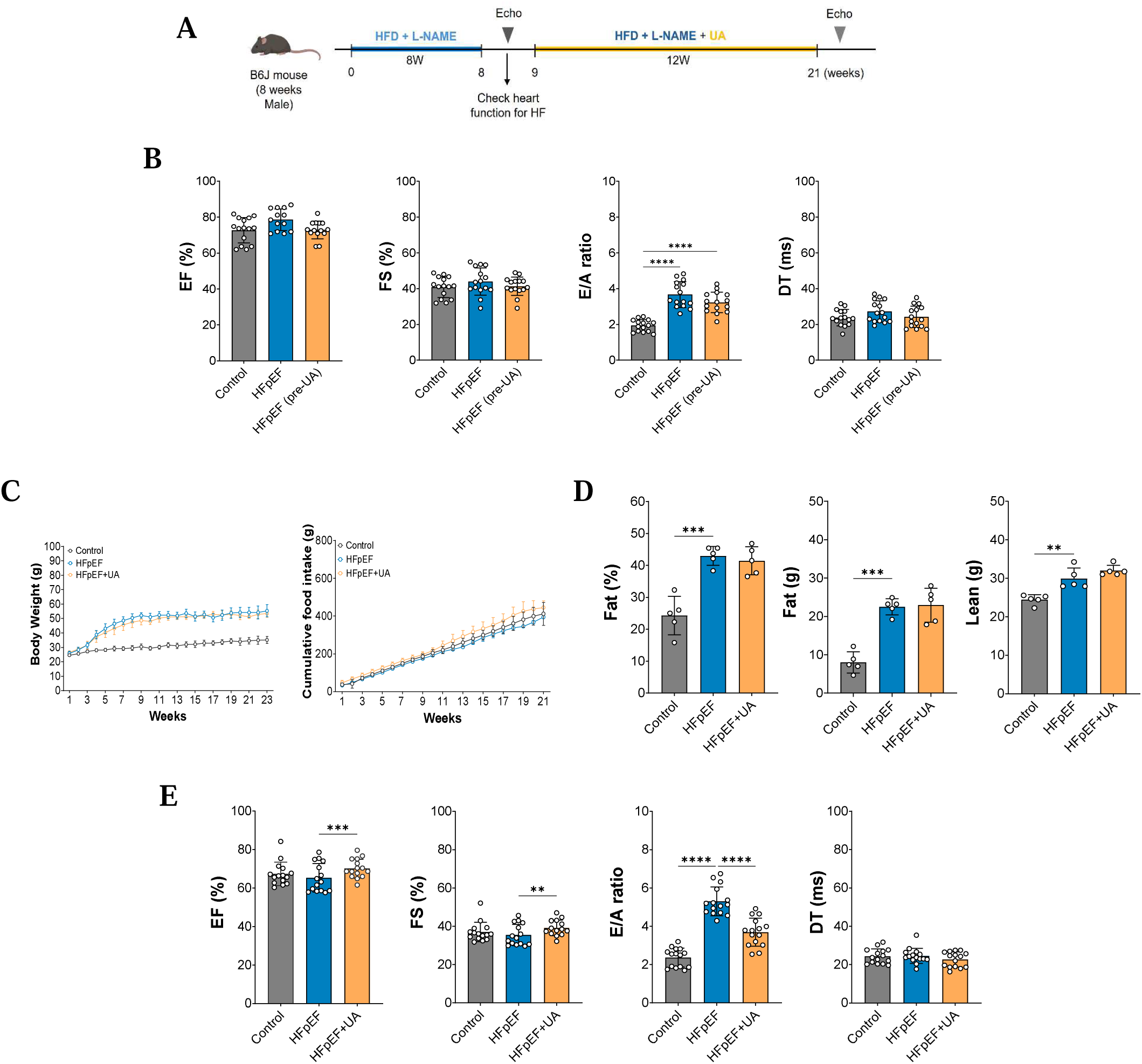
Experimental design and physiological characterization of a two-hit HFpEF mouse model with UA intervention. **A**, Schematic overview of the experimental timeline. Eight-week-old male C57BL/6J mice were subjected to a two-hit HFpEF protocol consisting of HFD and L-NAME administration for 8 weeks, followed by an additional 12 weeks of continued HFD and L-NAME with or without UA treatment. Echocardiographic assessments were performed at weeks 8 and 21. **B**, Echocardiographic evaluation of systolic and diastolic function at week 12, including ejection fraction (EF), fractional shortening (FS), E/A ratio, and deceleration time (DT). **C**, Longitudinal measurements of body weight (BW) and cumulative food intake over the study period in Control, HFpEF, and HFpEF+UA groups. **D**, Endpoint body composition analysis showing fat percentage, fat mass, and lean mass at study termination. **E**, Echocardiographic assessment at week 21 demonstrating systolic and diastolic functional parameters following UA treatment. All data are presented as mean ± SEM. Statistical significance was assessed using two-way ANOVA with Dunnett’s multiple comparisons test or Kruskal–Wallis test with Dunn’s post hoc test, as appropriate.

Terminal echocardiographic assessment at week 20 revealed significant functional improvements in the UA-treated mice. Both systolic parameters (ejection fraction and fractional shortening) were increased compared to those in untreated HFpEF mice. Furthermore, the diastolic function markedly improved, with the E/A ratio decreasing toward the control values (Fig. 1E), consistent with restored ventricular relaxation and filling dynamics. Collectively, these data demonstrated that UA treatment ameliorated cardiac dysfunction in HFpEF without confounding metabolic alterations.

### 3.2 UA treatment attenuated pathological cardiac hypertrophy and fibrosis in HFpEF mice

To examine the structural basis of diastolic dysfunction in HFpEF, we evaluated whether UA treatment could reverse the pathological remodeling at the organ and cellular levels. H&E staining of short-axis cardiac sections revealed concentric left ventricular hypertrophy with thickened walls in HFpEF mice (Fig. 2A). This was confirmed by gravimetric analysis; the heart weight-to-tibia length ratio increased in HFpEF mice compared to controls, and the effect was attenuated by UA treatment (Fig. 2B). HFpEF mice developed pulmonary congestion because of elevated filling pressures, as evidenced by the increased lung weight-to-tibia length ratio (Fig. 2C). UA treatment normalized this parameter, consistent with the improved E/A ratio observed on echocardiography (Fig. 1E).

**Figure 2.**
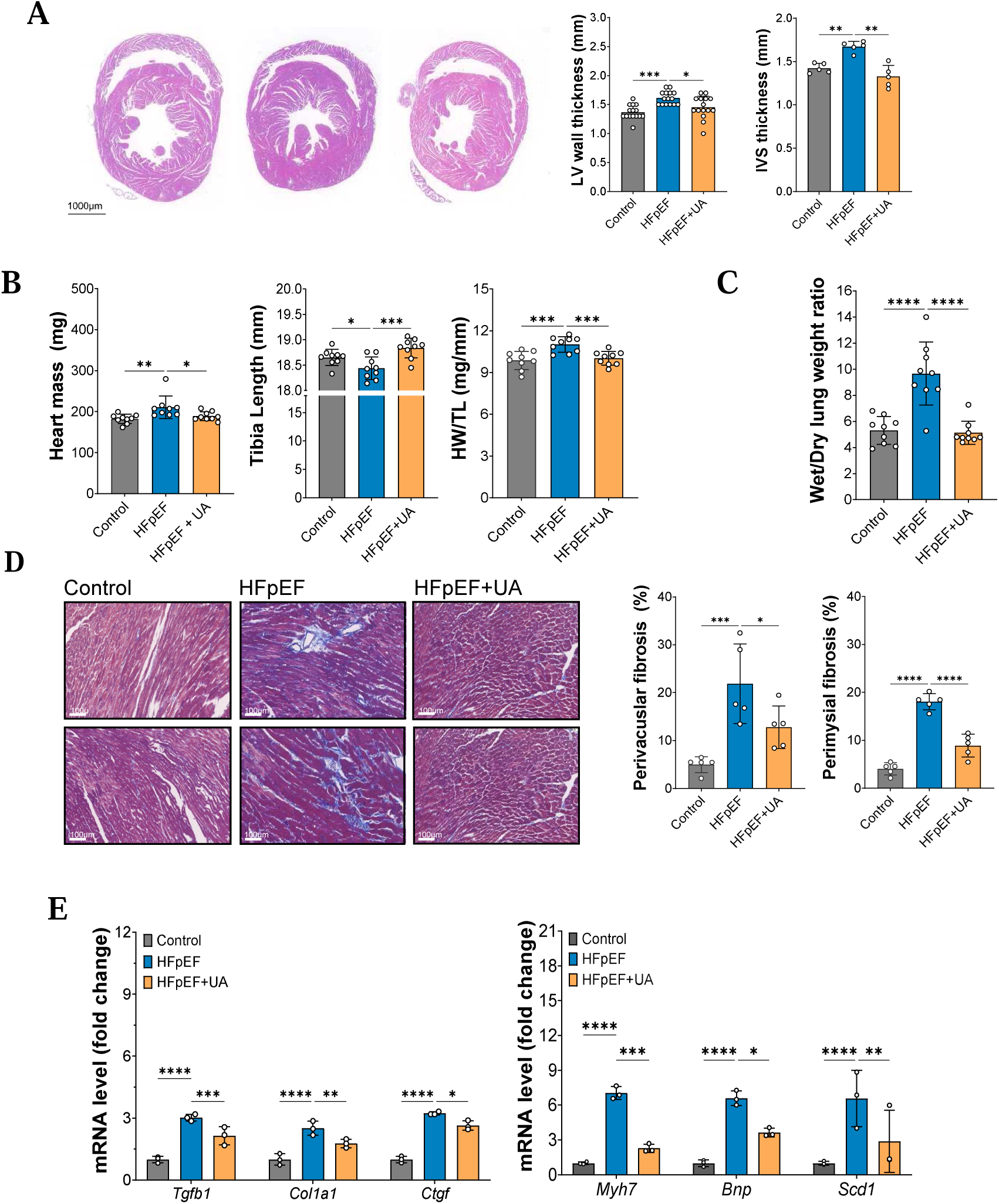
Structural and molecular characterization of cardiac remodeling in a two-hit HFpEF mouse model. **A**, Representative H&E stained short-axis cardiac sections from Control, HFpEF, and HFpEF+UA mice, shown from left to right. Quantification of left ventricular (LV) wall thickness and interventricular septal (IVS) thickness is shown on the right. Scale bar, 1000 µm. **B**, Gravimetric and anatomical measurements including heart mass, tibia length, and heart weight to tibia length ratio (HW/TL). **C**, Lung wet to dry weight ratio measured at study endpoint as an index of pulmonary congestion. **D**, Representative Masson’s trichrome (MT) stained LV sections illustrating myocardial collagen deposition across experimental groups. Quantification of perivascular and perimysial fibrosis is shown on the right. Scale bar, 100 µm. **E**, Quantitative PCR analysis of fibrosis associated genes (*Tgfb1*, *Col1a1*, and *Ctgf*) and hypertrophic/metabolic stress-related markers (*Myh7*, *Bnp*, and *Scd1*) in heart tissue. Gene expression levels were normalized to internal controls and expressed relative to Control. All Data are presented as mean ± SEM. Statistical significance was assessed using two-way ANOVA with Dunnett’s multiple comparisons test or Kruskal–Wallis test with Dunn’s post hoc test, as indicated. *P<0.05, **P<0.01, ***P<0.001.

Myocardial fibrosis, a hallmark of diastolic stiffness in HFpEF, was evaluated using Masson’s trichrome staining (Fig. 2D). HFpEF hearts exhibited extensive collagen deposition in both interstitial and perivascular compartments, whereas UA treatment markedly reduced the extracellular matrix accumulation (Fig. 2D). At the molecular level, fibrosis-associated genes were significantly upregulated in the HFpEF myocardium, including transforming growth factor beta 1 (*Tgfb1*), collagen type 1 alpha 1 (*Col1a1*), and connective tissue growth factor (*Ctgf*). UA treatment attenuated these transcriptional changes and restored the expression levels to control values (Fig. 2e). Similarly, the hypertrophic gene markers including myosin heavy chain 7 (*Myh7*) and natriuretic peptide B (*Nppb*) were elevated in the HFpEF group and normalized by UA treatment (Fig. 2E). These histological and molecular findings demonstrated that UA mitigated the maladaptive structural remodeling in HFpEF, thereby providing a mechanistic basis for the preservation of diastolic function.

### 3.3 UA treatment restored mitochondrial integrity and activated mitophagy through AMPK-mTOR signaling

Mitochondrial quality control is essential for maintaining cardiac bioenergetics under pathological stress^7^. Therefore, we examined whether the functional improvements conferred by UA were accompanied by the restoration of the mitochondrial ultrastructure and mitophagic flux. Figure 3a shows representative TEM images of mouse heart tissue. TEM imaging revealed pronounced mitochondrial damage in HFpEF hearts, including fragmentation, swelling, and loss of cristae density. UA treatment preserved mitochondrial architecture, maintained the cristae organization, and prevented vacuolization (Fig. 3A).

**Figure 3.**
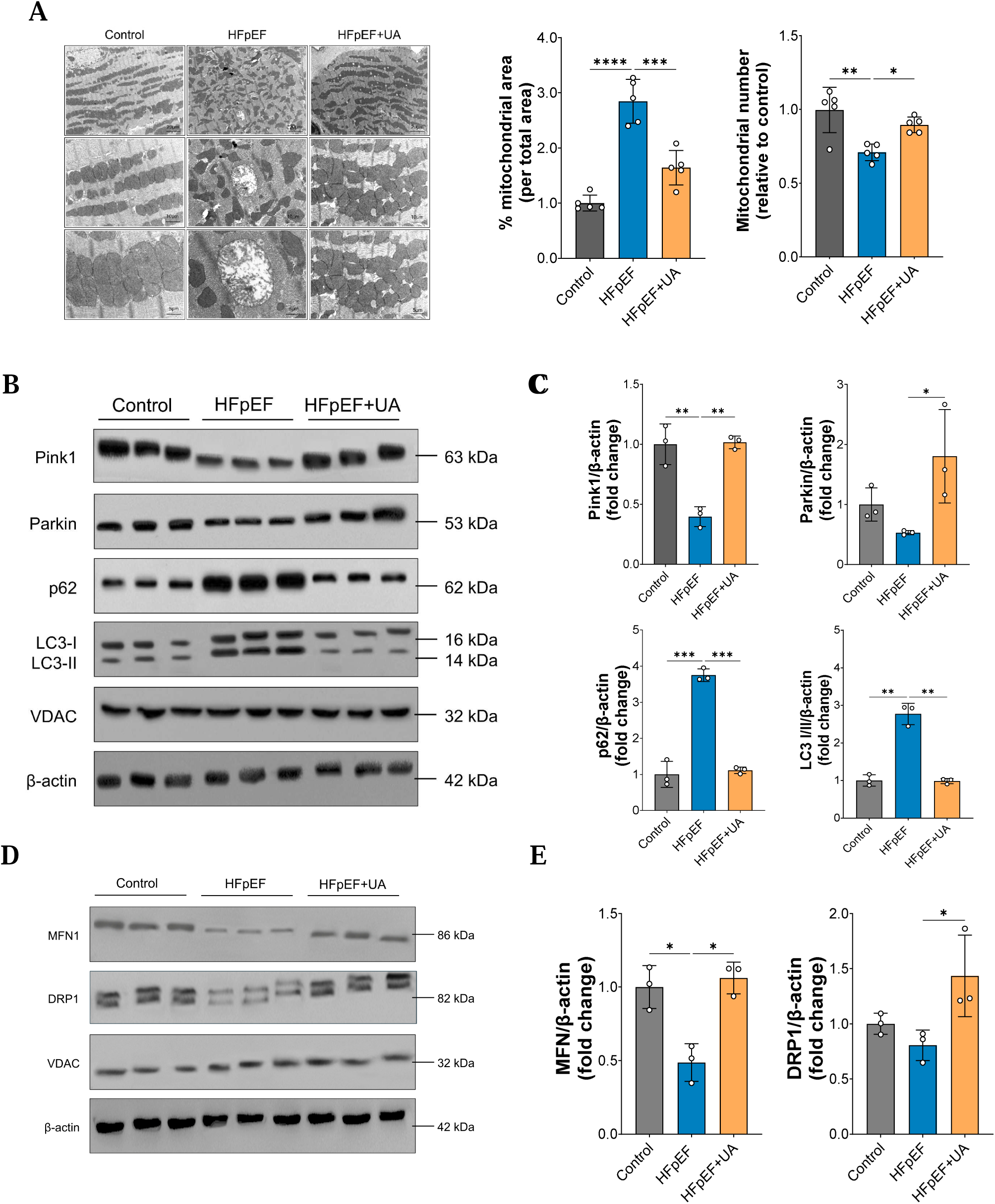
Mitochondrial ultrastructure and mitophagy-related signaling in HFpEF hearts with UA treatment. **A**, TEM images of heart tissue from Control, HFpEF, and HFpEF+UA mice. Quantification of mitochondrial area (percentage of total tissue area) and mitochondrial number normalized to Control is shown on the right. **B**, Immunoblot analysis of mitophagy-related proteins, including PINK1, Parkin, p62/SQSTM1, and LC3-I/II, in cardiac tissue lysates. **C**, Densitometric quantification of PINK1, Parkin, p62, and LC3-II levels normalized to β-actin. **D**, Immunoblot analysis of mitochondrial dynamics–associated proteins MFN1 (fusion) and DRP1 (fission). **E**, Quantification of MFN1 and DRP1 protein expression normalized to β-actin. Data are presented as mean ± SEM. Statistical significance was assessed using two-way ANOVA followed by Dunnett’s multiple comparisons test, as indicated. *P<0.05, **P<0.01, ***P<0.001.

To assess whether these structural improvements reflected the enhanced clearance of damaged organelles, we evaluated the PINK1-Parkin-mediated mitophagy pathway activated by UA^45^. Immunoblotting revealed diminished expression of both PINK1 and Parkin in the HFpEF myocardium (Figs. 3B, C). Markers of autophagic flux, the LC3-II/LC3-I ratio and p62/SQSTM1, indicated impaired autophagy in HFpEF (Figs. 3B, C). UA administration comprehensively restored autophagy by increasing PINK1 and Parkin expression, enhancing LC3-II conversion, and promoting efficient p62 clearance. Consistent with these findings, mitophagy regulatory genes, including *Rab7* and *Pink1*, were transcriptionally downregulated in HFpEF and restored by UA treatment (Supplementary Fig. 1A). In addition to defective mitophagy, HFpEF hearts exhibited abnormalities in mitochondrial dynamics. Immunoblot analyses showed reduced expression of the fusion protein MFN1 and increased levels of the fission regulator DRP1, indicating a fragmented mitochondrial network (Figs. 3D, E). UA treatment restored MFN1 abundance and normalized DRP1 expression, suggesting that UA not only reactivated mitophagy but also rebalanced the mitochondrial fusion-fission dynamics to support mitochondrial integrity.

Next, we investigated the upstream signaling mechanisms underlying mitophagy reactivation. AMP-activated protein kinase (AMPK), a master regulator of cellular energy homeostasis, activates mitophagy, whereas the mechanistic target of rapamycin (mTOR) suppresses it^46,47^. HFpEF hearts showed decreased AMPK phosphorylation but increased mTOR phosphorylation (Supplementary Figs. 1A, C). UA treatment reversed this maladaptive signaling profile by increasing p-AMPK levels and suppressing p-mTOR. Consistent with AMPK activation, UA treatment restored the phosphorylation of ULK1 and Beclin-1, which are direct AMPK targets that are critical for autophagosome initiation (Supplementary Fig. 1C). These findings demonstrated that UA treatment alleviated mitochondrial dysfunction in HFpEF by reactivating AMPK-dependent mitophagy.

Next, we validated these in vivo findings using cultured cardiomyocytes. The mt-Keima reporter was used in H9c2 cells to directly visualize the mitophagic flux^48^. When exposed to HFpEF-like stress (palmitate+L-NAME), cells predominantly displayed green fluorescence, indicating suppressed mitophagy. Treatment with UA markedly increased the red fluorescence, which increased the red-to-green ratio and demonstrated restored mitochondrial delivery to lysosomes (Supplementary Figs. 2A, B). We then assessed the mitochondrial function using a Seahorse XF analyzer. HFpEF stress reduced the basal oxygen consumption, ATP-linked respiration, and maximal respiratory capacity. However, UA treatment reversed these deficits, restoring the basal and maximal OCR and recovering the spare respiratory capacity, i.e., the energetic reserve available to meet the increased demand (Supplementary Fig. 2C). Finally, we confirmed that UA required functional lysosomes using bafilomycin A1 (BafA1), which prevents autophagosome-lysosome fusion^49^. Without BafA1, UA treatment increased the LC3-II conversion while clearing p62, which was consistent with the productive flux (Supplementary Figs. 2D, E). However, BafA1 co-treatment caused the accumulation of LC3-II to high levels while completely preventing p62 clearance. This blockade proved that UA operated through the canonical autophagy-lysosome pathway (Supplementary Figs. 2D, E). Collectively, these cellular assays confirmed that UA treatment activated mitophagy, eliminated dysfunctional mitochondria, and preserved respiratory capacity.

### 3.4 UA treatment remodeled the gut microbiome and reduced the systemic ceramide burden

Accumulating evidence has implicated gut dysbiosis in the pathogenesis of HFpEF through the production of cardiotoxic metabolites^50,51^. Previous studies have reported that UA treatment can restore gut dysbiosis by modulating the intestinal microbiota and its functional profile in response to metabolic stress^52,53^. To determine whether the therapeutic effects of UA involve the modulation of the gut microbiome in HFpEF mice, we performed shotgun metagenomic sequencing of fecal samples. The alpha diversity, which was assessed using the Shannon index, was reduced in HFpEF but significantly improved by UA treatment (Fig. 4A). Principal coordinate analysis based on the Bray-Curtis dissimilarity revealed a distinct clustering of gut microbial communities across the groups (Fig. 4A and Supplementary Fig. 3A). The Firmicutes/Bacteroidetes ratio—a marker of metabolic dysbiosis^54^—was elevated in HFpEF and normalized by UA treatment (Supplementary Fig. 3B). At the taxonomic level, HFpEF was characterized by the selective expansion of genera previously linked to ceramide synthesis, including *Bacteroides*, *Parabacteroides*, *Alistipes*, and *Phocaeicola*. All four taxa exhibited a significantly increased relative abundance in HFpEF mice, consistent with their reported roles in promoting ceramide synthesis (Fig. 4B and Supplementary Fig. 3C)^55–57^. Notably, UA treatment suppressed the overgrowth of these ceramide-associated genera, restoring their abundance to control levels (Fig 4B).

**Figure 4.**
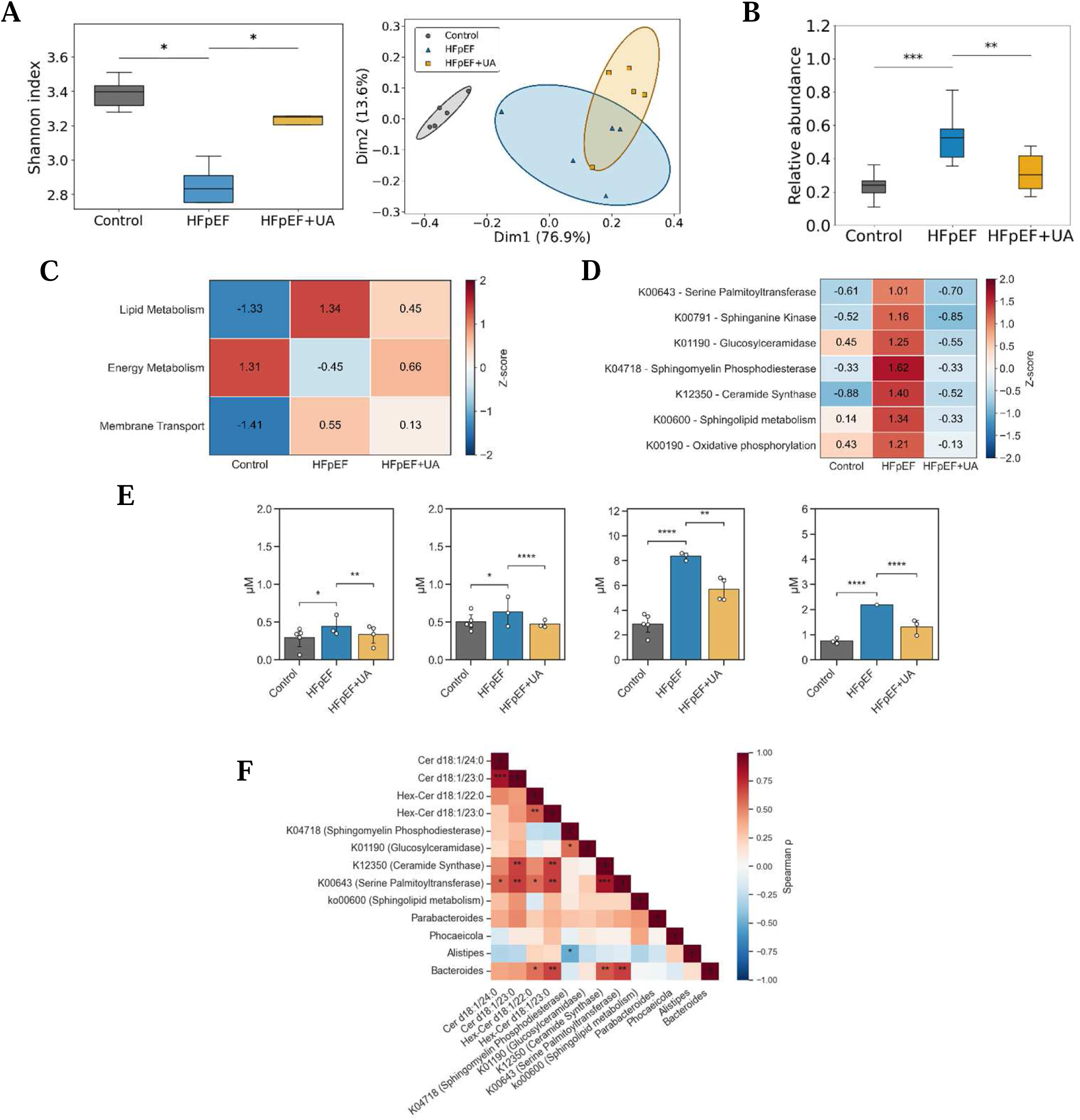
Gut microbiome composition, functional profiling, and ceramide-related lipid alterations in HFpEF with UA treatment. **A**, Microbial diversity analyses showing Shannon α-diversity and Bray-Curtis–based principal coordinates analysis (PCoA) of fecal metagenomic profiles from Control, HFpEF, and HFpEF+UA mice. **B**, Relative abundance of microbial taxa implicated in sphingolipid (ceramide) biosynthesis across experimental groups. **C**, Z-score heatmap of major KEGG functional categories derived from metagenomic functional profiling. **D**, Z-score heatmap of KEGG orthology modules related to ceramide and sphingolipid metabolism. **E**, Targeted plasma lipidomic quantification of selected ceramide and hexosyl-ceramide species, shown from left to right as Cer d18:1/23:0, Cer d18:1/24:0, Hex-Cer d18:1/22:0, and Hex-Cer d18:1/23:0. **F**, Spearman correlation matrix integrating dominant microbial taxa, ceramide-associated KEGG modules, and circulating ceramide species. Data are presented as mean ± SEM. Statistical significance was assessed using two-way ANOVA followed by Dunnett’s multiple comparisons test. *P<0.05, **P<0.01, ***P<0.001, ****P<0.0001.

To assess the functional consequences, we analyzed the microbial metabolic capacity using KEGG orthology enrichment. The HFpEF microbiomes displayed significant enrichment of lipid metabolism pathways, particularly sphingolipid biosynthesis (Figs. 4C, D). Specifically, genes encoding ceramide biosynthetic enzymes such as serine palmitoyltransferase (K00643), sphingomyelin phosphodiesterase (K04718), and ceramide synthase (K12350) were upregulated in HFpEF and suppressed by UA treatment (Fig. 4D). These microbial alterations resulted in systemic metabolic changes. Targeted plasma lipidomics revealed that HFpEF mice exhibited elevated levels of ceramides (Cer d18:1/23:0 and Cer d18:1/24:0) and hexosylceramides (HexCer d18:1/22:0 and HexCer d18:1/23:0), which are lipid species that are strongly associated with cardiovascular disease risk (Fig. 4E and Supplementary Fig. 3D)^58,59^. UA treatment significantly reduced the levels of these ceramides in circulation. Integrative correlation analysis revealed positive correlations among the abundance of microbiota, expression of genes involved in ceramide synthesis, and plasma ceramide concentrations (Fig. 4F). These findings suggested that UA treatment remodeled the gut microbiome to suppress microbial sphingolipid biosynthesis. This reduced the systemic ceramide levels and may contribute to the cardioprotective effects of UA in patients with HFpEF.

### 3.5 UA treatment prevented maladaptive metabolic reprogramming in human cardiomyocytes under HFpEF-like stress

To validate these findings in a human system, we exposed human iPSC-derived cardiomyocytes (hiPSC-CMs) to HFpEF-like stress (palmitate+L-NAME) (Fig. 5A). The combined stress significantly induced changes in inflammation, cardiac fibrosis, and hypertrophy. These changes were attenuated by co-treatment with UA (Fig. 5B). To resolve the cellular heterogeneity and transcriptional remodeling at a single-cell resolution, we performed snRNA-seq on hiPSC-CMs under control, HFpEF-like stress (HL), and UA-treated conditions. Unsupervised clustering and UMAP dimensionality reduction revealed distinct transcriptional states (Fig. 5C). Cell type annotation revealed ten subclusters, including ventricular cardiomyocytes, as well as atrial-like, proliferative, and stressed subclusters (Fig. 5C). Notably, HL stress induced the emergence of an aberrant hepatocyte-like cellular subpopulation characterized by the expression of genes typically related to hepatocyte features, including albumin (*ALB*), transthyretin (*TTR*), and apolipoprotein A1 (*APOA1*)^60,61^.

**Figure 5.**
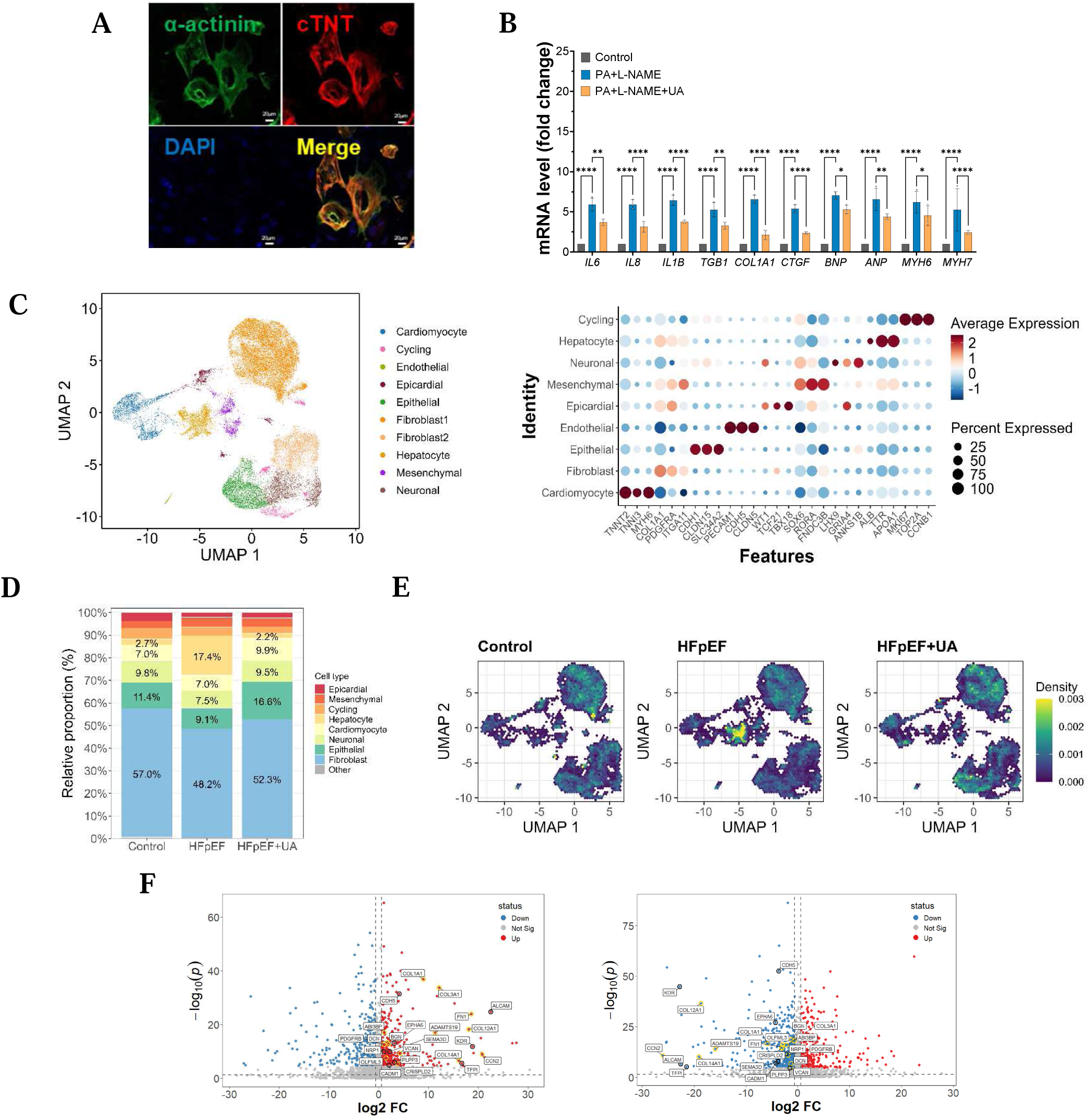
Transcriptional and cellular state profiling of human cardiomyocytes under HFpEF-like stress with urolithin A treatment. **A**, Immunofluorescence staining of human induced pluripotent stem cell–derived cardiomyocytes (hiPSC-CMs) for α-actinin, cardiac troponin T (cTnT), and DAPI, confirming sarcomeric organization and cardiomyocyte identity. **B**, Quantitative PCR analysis of inflammatory, fibrotic, and hypertrophic marker genes in Control, HFpEF-like stress (palmitate+L-NAME; HL), and HL+UA conditions. **C**, snRNA-seq–based cell-state mapping of hiPSC-derived cardiomyocytes (hiPSC-CMs). UMAP visualization and dot plots of canonical marker genes were used to annotate major cardiomyocyte and non-cardiomyocyte–like transcriptional subclusters across experimental groups. **D**, Relative proportions of cardiomyocyte subclusters and hepatocyte-like populations across Control, HL, and HL+UA conditions. **E**, UMAP density plots depicting transcriptomic distribution patterns under each experimental condition, shown from left to right as Control, HFpEF-like stress (HL), and HL+UA. **F**, Volcano plots of differentially expressed genes within the hepatocyte-like subcluster comparing HL versus Control and HL+UA versus HL.

This “hepatocyte-like” cluster was expanded under HL conditions; however, UA treatment substantially reduced this population to control proportions (Figs. 5D, E). Differential gene expression (DGE) analysis within this maladaptive cluster revealed the coordinated upregulation of profibrotic and proinflammatory transcripts under HL stress (Fig. 5F). Treatment with UA suppressed these pathological signatures. These findings demonstrated that HFpEF-induced stress triggered maladaptive phenotypic plasticity in human cardiomyocytes, which manifested as the acquisition of profibrotic and proinflammatory transcriptional states. UA prevented this detrimental reprogramming and preserved the cardiomyocyte identity and function.

### 3.6 UA treatment prevented the cardiomyocyte fibrogenic transition and restored mitochondrial quality control

To elucidate the transcriptional mechanisms underlying the cardioprotective effects of UA at a single-cell resolution, we performed subcluster analysis and trajectory inference on the cardiomyocyte population from our snRNA-seq dataset. The DGE analysis of the cardiomyocyte population revealed that UA treatment completely reversed the HFpEF-induced transcriptional signature (Fig. 6A). UA treatment downregulated a broad panel of profibrotic genes and simultaneously upregulated mitophagy regulators, which were suppressed under stress conditions (Fig. 6B). To map the continuum of cardiomyocyte transcriptional states, we employed pseudotime trajectory analysis^62^. This approach resolved distinct subpopulations, including ventricular, immature atrial, and conduction-type cardiomyocytes, along with a population highly enriched for extracellular matrix (ECM) gene expression, designated the “Fibro_core” cluster (Figs. 6C, D and Supplementary Fig. 4A). Quantification of cluster proportions revealed that HFpEF stress markedly expanded the Fibro_core population, and UA treatment effectively reduced this population to below control levels (Supplementary Fig. 4B), indicating not only prevention but also active suppression of fibrogenic reprogramming.

**Figure 6.**
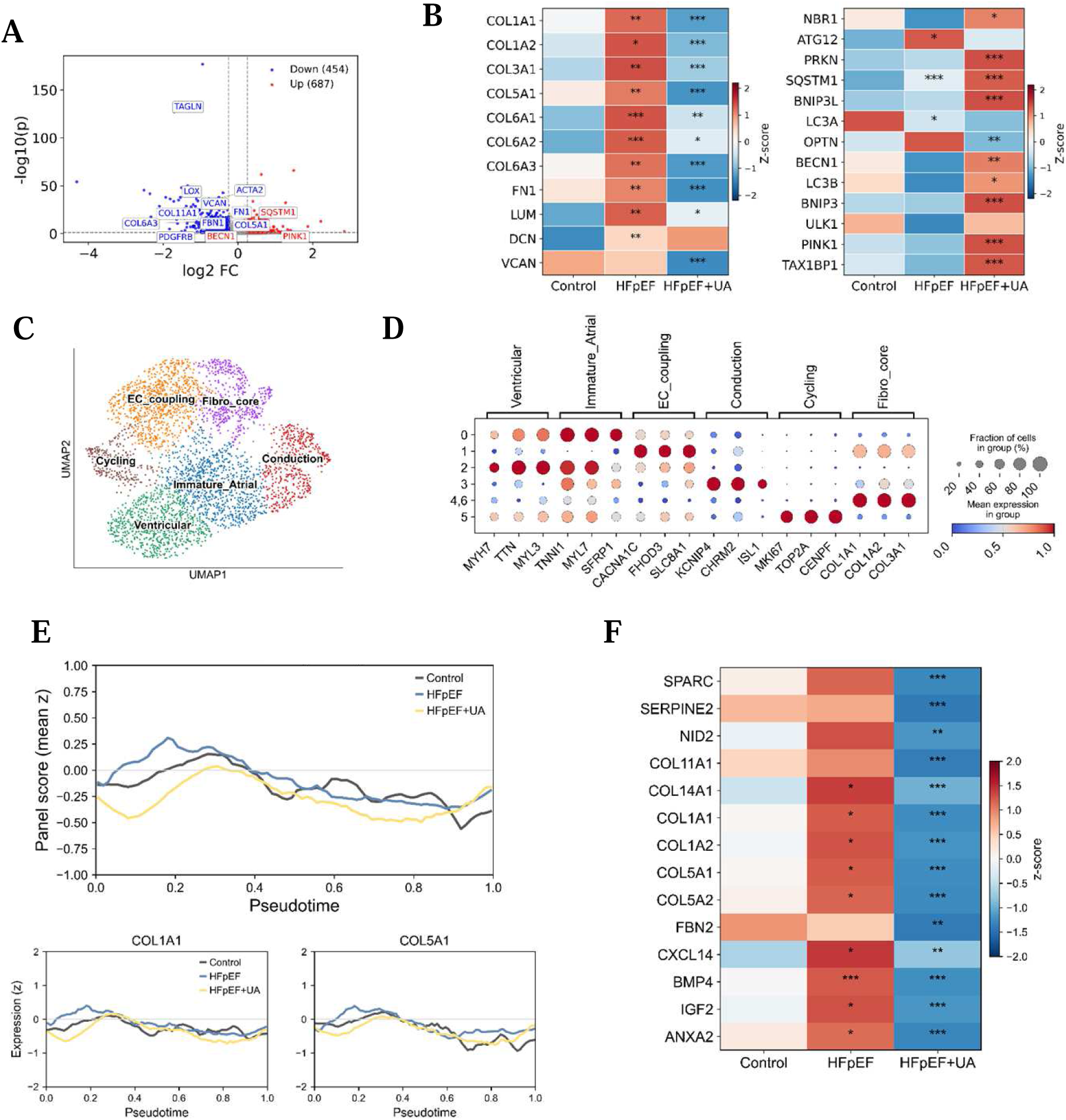
Transcriptional programs related to fibrosis and mitochondrial quality control in HFpEF cardiomyocytes with UA treatment. **A**, Volcano plot of differentially expressed genes comparing HFpEF+UA and HFpEF cardiomyocytes, highlighting transcripts significantly upregulated or downregulated following UA treatment. **B**, Z-score heatmaps of extracellular matrix (ECM)/collagen genes (left) and mitophagy-related genes (right) across Control, HFpEF, and HFpEF+UA groups. Statistical significance is indicated as *P<0.05, **P<0.01, ***P<0.001. **C**, UMAP embedding of cardiomyocyte nuclei delineating major transcriptional modules, including Ventricular, Immature_Atrial, Conduction, EC_coupling, Cycling, and ECM-enriched Fibro_core populations. **D**, Dot plot of canonical marker genes used to define each transcriptional module, with dot size indicating the fraction of expressing cells and color representing scaled mean expression. **E**, Pseudotime trajectories of representative genes from a curated fibrosis core 14-gene module, including COL1A1 (left) and COL5A1 (right), showing dynamic expression patterns along the cardiomyocyte state continuum. **F**, Heatmap of fibrosis-associated genes across pseudotime, illustrating group-specific expression patterns in Control, HFpEF, and HFpEF+UA cardiomyocytes. Data are presented as mean±SEM.

Analysis of gene expression dynamics along the pseudotime trajectory revealed that profibrotic genes were progressively induced during the early to-mid trajectory phases in HFpEF cardiomyocytes (Fig. 6E and Supplementary Fig. 4C). Quantitative scoring of the fibrosis gene module confirmed that HFpEF significantly elevated fibrosis scores during these critical transition phases, whereas UA treatment maintained low scores compared to controls (Fig. 6E and Supplementary Fig. 4D). Within the Fibro_core cluster, DGE analysis showed that UA treatment markedly suppressed key drivers of ECM remodeling and fibrosis, including secreted protein acidic and rich in cysteine (*SPARC*), serpin family E member 2 (*SERPINE2*), *COL1A1*, and bone morphogenetic protein 4 (*BMP4*)(Fig. 6F)^63–65^. These findings demonstrated that UA treatment prevented the pathological phenotypic plasticity of human cardiomyocytes by blocking their trajectory toward profibrotic transcriptional states while concurrently reactivating mitochondrial quality control programs.

## DISCUSSION

Cardiovascular diseases remain the leading global cause of morbidity and mortality, with HF representing a growing burden^66^. HFpEF represents a major therapeutic challenge largely because it is not an isolated cardiac condition but a heterogeneous syndrome driven by multiple systemic comorbidities, including obesity, hypertension, diabetes, chronic kidney disease, and atrial fibrillation^67^. Accumulating evidence indicates that mitochondrial dysfunction is a central mechanistic link between systemic metabolic perturbations and cardiomyocyte pathology in HFpEF^6,68^. Therapeutic strategies targeting MQC may address the fundamental defects in myocardial energetics and contractile function that underlie this syndrome^69^.

We demonstrated that UA administration ameliorated HFpEF pathology in a two-hit mouse model. UA treatment improved the cardiac structure and function, reduced the left ventricular mass, improved diastolic parameters, and decreased interstitial fibrosis and pulmonary congestion (Fig. 2). We also found that UA treatment restored the phosphorylation of AMPK and its downstream targets (ULK1 and Beclin-1) while suppressing the elevated mTOR activity observed in HFpEF hearts. This signaling shift re-established the PINK1-Parkin-mediated mitophagic flux, which promoted the clearance of damaged mitochondria and preserved the cristae architecture (Fig. 3 and Supplementary Figs. 1 and 2). These findings extend previous observations of the beneficial effects of UA in skeletal muscles and aging models^11,45^, suggesting that its mitophagy-enhancing properties are operational in the stressed myocardium.

A notable finding of our study was the systemic impact of UA on the gut-heart metabolic axis. UA treatment modulated the gut microbiome composition and reduced the abundance of genera, including *Bacteroides* and *Parabacteroides* (Fig. 4 and Supplementary Fig. 3). Metagenomic analysis revealed the downregulation of bacterial genes encoding enzymes involved in de novo ceramide biosynthesis. This microbial shift was correlated with reduced circulating ceramide levels. Given that ceramides promote mitochondrial dysfunction, inflammasome activation, and fibroblast differentiation^70,71^, a UA-induced decline in systemic ceramides may interrupt a lipid-driven pathogenic feedback loop, complementing the direct myocardial effects. Our snRNA-seq analysis of human iPSC-CMs provided insights into the cellular responses to metabolic stress and UA treatment. Under HFpEF-like conditions, UA treatment attenuated pathological transcriptional programs, reducing the expression of profibrotic markers while maintaining the expression of genes associated with normal cardiomyocyte function and MQC (Figs. 5 and 6). These observations suggest that preserving mitochondrial health may be important for maintaining cardiomyocyte transcriptional identity, although the directional relationship between mitochondrial dysfunction and transcriptional reprogramming remains unclear.

Our study had several limitations. First, although our two-hit model recapitulated the key features of HFpEF, it represented a relatively short-term intervention in young mice and may not have fully captured the chronic age-related pathophysiology of human HFpEF. Second, although we demonstrated strong correlations among microbiome composition, ceramide levels, and cardiac phenotypes, establishing definitive causal relationships requires additional experimental approaches, such as germ-free models or targeted microbial manipulations. Third, while human iPSC-CMs provide human-relevant mechanistic insights, they exhibit a relatively immature phenotype compared to the adult myocardium, potentially limiting translatability.

Despite these limitations, our study provided mechanistic evidence that UA targets multiple interconnected pathways involved in the pathogenesis of HFpEF. By restoring MQC, modulating the gut microbiome composition and systemic ceramide metabolism, and preventing maladaptive cardiomyocyte transcriptional changes, UA addressed several core features of HFpEF pathophysiology in this preclinical model. These findings provide a rationale for the clinical investigation of UA as a potential therapeutic approach for HFpEF, although substantial additional work is required to establish its safety, efficacy, and optimal dosing in humans.

## Acknowledgements

None

## Author Contributions

C.-M.O. conceived the study. C.-M.O., and S.-W.P. designed the study. H.S. performed most experiments with help from C.Y., Y.K., J.K., J.Y.L., and D.R.., Y.J.C. and W.J T. Ziv helped in the design, investigation, and analysis of transriptomics C.-M.O. and H.S. drafted the article with input from all authors.

## Supplemental Material

Supplemental Methods

Table S1

Figures S1-S4

Major Resources Table

## Abbreviations

AMPK: AMP-activated protein kinase
ATCC: American Type Culture Collection
BafA1: Bafilomycin A1
BMP4: Bone morphogenetic protein 4
BNP / Nppb: B-type natriuretic peptide
BW: Body weight
CCCP: Carbonyl cyanide m-chlorophenyl hydrazone
Cer: Ceramide
CTGF / Ctgf: Connective tissue growth factor
DXA: Dual-energy X-ray absorptiometry
ECM: Extracellular matrix
EF: Ejection fraction
FBS: Fetal bovine serum
FS: Fractional shortening
GEO: Gene Expression Omnibus
H&E: Hematoxylin and eosin
HF: Heart failure
HFpEF: Heart failure with preserved ejection fraction
HFD: High-fat diet
HL: HFpEF-like stress (palmitate + L-NAME)
hiPSC-CMs: Human induced pluripotent stem cell–derived cardiomyocytes
HUMAnN: The HMP Unified Metabolic Analysis Network
IVS: Interventricular septum
KEGG: Kyoto Encyclopedia of Genes and Genomes
LC3: Microtubule-associated protein 1A/1B-light chain 3
L-NAME: Nω-nitro-L-arginine methyl ester
LV: Left ventricle / Left ventricular
MQC: Mitochondrial quality control
mt-Keima: Mitochondria-targeted Keima fluorescent reporter
mTOR: Mechanistic target of rapamycin
OCR: Oxygen consumption rate
PCoA: Principal coordinates analysis
PINK1: PTEN-induced putative kinase 1
PVDF: Polyvinylidene difluoride
qPCR / qRT-PCR: Quantitative real-time polymerase chain reaction
RNA-seq: RNA sequencing
SEM: Standard error of the mean
snRNA-seq: Single-nucleus RNA sequencing
SRA: Sequence Read Archive
TEM: Transmission electron microscopy
Tgfb1: Transforming growth factor beta 1
UA: Urolithin A
UMAP: Uniform Manifold Approximation and Projection

